# Different macroevolutionary routes to becoming a biodiversity hotspot

**DOI:** 10.1101/452128

**Authors:** J. Igea, A. J. Tanentzap

## Abstract

Why is species diversity so unevenly distributed across different regions on Earth? Regional differences in biodiversity may stem from differences in rates of speciation and dispersal and colonization times, but these hypotheses have rarely been tested simultaneously at a global scale. Here we uncovered the routes that generated hotpots of mammal and bird biodiversity by analyzing the tempo and mode of diversification and dispersal within major biogeographic realms. Hotspots in tropical realms had higher rates of speciation whereas those in temperate realms received more immigrant species from their surrounding regions. We also found that hotspots had higher spatial complexity and energy availability, providing a link between the environment and macroevolutionary history. Our study highlights how assessing differences in macroevolutionary history can help to explain why biodiversity varies so much worldwide.

## Main Text

Biodiversity is extremely unevenly distributed across the globe and understanding why has long fascinated biologists (*1*). For example, there are many exceptions to the tendency for species richness to increase towards the Equator - widely studied as the latitudinal diversity gradient (*1–3*). This finer-scale association between biodiversity and geography (*4*) is exemplified by the 35 terrestrial biodiversity hotspots proposed by Myers *et al*. (*5, 6*) for conservation purposes based on plant endemicity and habitat loss. A third of Myers’ hotspots were located in temperate zones and were more diverse than many regions closer to the Equator, demonstrating that high levels of species richness can also be found outside the tropics.

Regional differences in biodiversity may ultimately arise through at least one of three macroevolutionary routes. First, differences in historic rates of *in situ* diversification (i.e., speciation minus extinction) can result in more species accumulating in some areas than others. Second, differences in historic rates of lineage dispersal can result in some areas acting as sources of species that are exported elsewhere and some that are sinks that import species (*7*). Finally, an older age of colonization of a region may promote diversity if there was more time to accumulate species, generating “museums” of biodiversity (*8*), as formalized in the “time-for speciation” hypothesis (*9, 10*). However, most studies of regional diversity patterns have not compared the relative importance of these different potential routes (*3*).

The three macroevolutionary routes that give rise to regional differences in biodiversity are at least partially paved by the environment (*1*). Some environmental variables may favor cladogenesis, such as past tectonic movements that generate isolation (*11*), whilst others may favor the establishment of immigrant species, such as historically stable climates that create regional refuges for species during periods of global change (*12*) and favor dispersal from environmentally similar areas, such as because of niche conservatism (*13*). Other environmental variables may favor both cladogenesis and immigration. For example, higher environmental energy might promote speciation by increasing mutation rates and shortening generation times (*14*), and also allow regions to hold more species by expanding their carrying capacity (*15*). Similarly, a higher speciation rate and local carrying capacity are both associated with physiographic heterogeneity (*16, 17*) and habitat complexity (*18, 19*).

Here we use terrestrial hotspots of mammal and bird biodiversity to understand how different macroevolutionary routes (*20*) generate extreme spatial differences in species diversity. We delineated hotspots using the number of species in an area divided by the inverse of range of those species. Hotspots based on this measure - known as weighted endemicity (WE, ref. *21*) and not to be confused with counting endemic species - identify the contribution of each area to global biodiversity more accurately than species richness, because widespread species are not counted in every area where they occur and so do not have a disproportionate influence on the metric. Therefore, hotspots based on WE will be more representative of the distribution of biodiversity across multiple regions on Earth than hotspots based on species richness, which are exclusively centered in the tropics. Using diversification rate and historical biogeography inference methods, we then tested which macroevolutionary routes could better explain the existence of mammal and bird hotspots across different regions on Earth. Efforts to reconstruct explicitly the historical rates of migration and diversification of biodiversity hotspots have largely focused on small clades or specific geographic regions (*20, 22, 23*), without a broader global context.

We first used global species maps (*24, 25*) to delineate hotspots. After overlaying species ranges with a grid of 100×100 km cells, we defined separate mammal and bird hotspots for subsequent analyses as the richest 20% of cells in terms of WE (Fig. S1). We chose this threshold to obtain clade-specific hotspots that were roughly equivalent in size to Myers’ hotspots (*5, 6*). Despite being partially delineated using habitat threat, we used Myers *et al*. (5) as a basis for comparison because they identified large spatial unevenness in biodiversity. They observed that 20% of the global land area was sufficient to retain many of the most biodiverse biomes on the planet (e.g., Andes, Sundaland, Madagascar, Mediterranean Basin) and >40% of all vertebrate species (*6*). Our resulting WE-based hotspots were largely overlapping with both Myers’ hotspots and hotspots of total species richness (see Methods), suggesting that all the measures captured a similar biological pattern.

Using hotspots and neighboring regions within six biogeographic realms, we assessed the spatial variation in the accumulation of ancient and recent lineages. To do this, we first obtained species-specific rates of diversification by estimating the Diversification Rate (DR) metric (*26*) for the most comprehensive phylogenies of mammals and birds (see Methods). DR captures the number of historic diversification events that lead to a given species, weighted by the relative age of those events, but does not explicitly model extinction (*26, 27*). We then built linear regression models that predicted the richness of species that were either ancient (i.e., older, with DR values in the 1^st^ quartile of the distribution) or recent (younger, with DR values in the 4^th^ quartile of the distribution). Total species richness in each cell (*27*) was the sole model predictor. By examining the residuals of these linear models, we determined which cells had an excess or a deficit of species in each of the two quartiles within each biogeographic realm (*27*). Our results revealed that hotspots generally had a deficit of ancient lineages and an excess of recent lineages when compared to neighboring regions, except in birds where evidence of the latter was mixed (Fig. 1). We confirmed that these results were robust both to phylogenetic uncertainty and the method used for estimating diversification rates (Figs S2-S3, Table S1, see Methods).

**Fig. 1.**
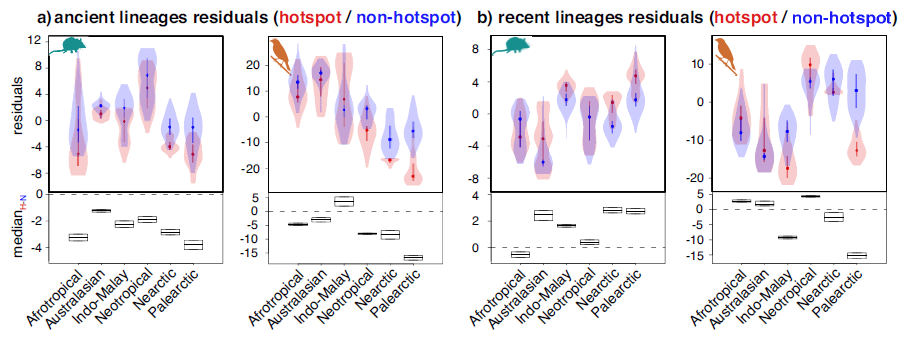
Hotspots are poor in ancient lineages and sometimes rich in recent lineages. Residuals from linear models predicting cell-specific richness of a) ancient and b) recent lineages in hotspots (H, shown in red) and non-hotspot regions (N, shown in blue). Positive residuals indicate a regional excess of ancient/recent lineages and negative residuals indicate a deficit. The lower panels show the median of the difference between a random hotspot point (H) and a random non-hotspot point (N) and the 95% confidence interval around that median calculated with a Wilcoxon rank sum test.

Next, we assessed whether differences in macroevolutionary routes generated the different patterns of accumulation of ancient and recent species in hotspot and non-hotspot regions across biogeographic realms. We found that the general deficit of ancient lineages and more variable excess of recent lineages in the hotspots compared to nearby regions resulted from contrasting macroevolutionary histories across biogeographic realms. We reached this conclusion by reconstructing assembly dynamics within hotspots and non-hotspots of each realm using historical biogeographic inference (*28*). We found that both mammal and bird species were generated at faster rates in the last 25 million years (Ma) within hotspots compared with nearby regions of largely tropical realms like Australasia, Indo-Malay and the Neotropics. By contrast, cladogenetic rates were similar or lower than surrounding areas in hotspots of the Afrotropics and temperate Palearctic and Nearctic realms (Fig. 2). Therefore, *in situ* cladogenesis could explain the accumulation of biodiversity in most tropical but not temperate hotspots. In temperate but not tropical realms, greater rates of historical dispersal rather than *in situ* cladogenesis could explain the accumulation of mammal and bird diversity within hotspots (Fig. 3). We found no evidence that colonization age alone could explain the differences in biodiversity as hotspots were generally colonized later than non-hotspot regions in temperate realms, and there was no consistent difference in the age of colonization across tropical realms (Fig. S4).

**Fig. 2.**
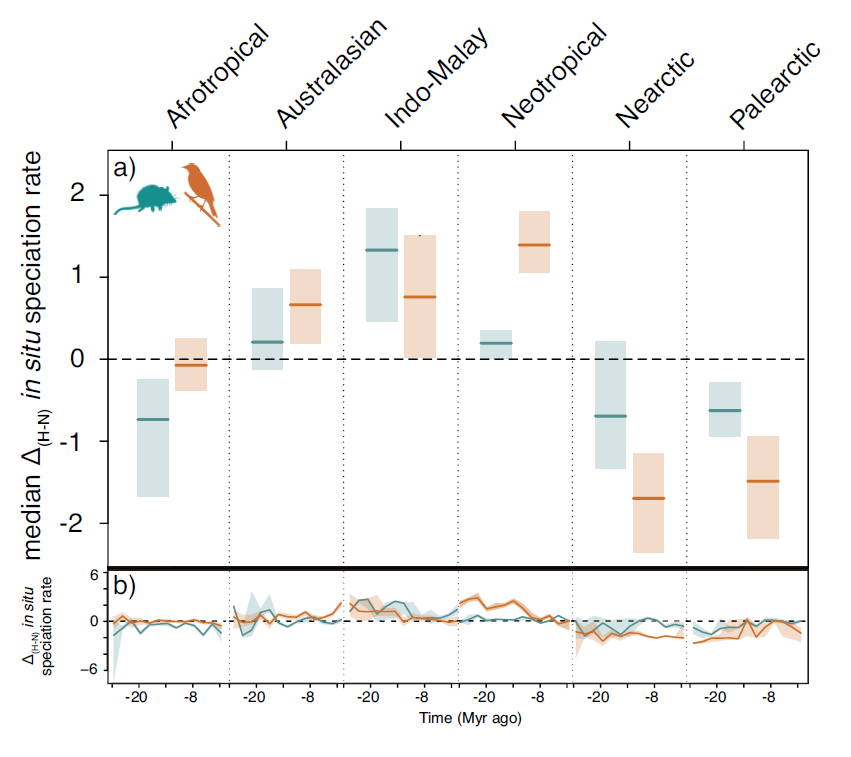
Contrasting rates of *in situ* cladogenesis in hotspots compared to surrounding non-hotspot regions. a) *In situ* cladogenesis rates between 2 to 26 Ma ago within non-hotspots were subtracted from rates within hotspots in each of six biogeographic realms and divided by the overall standard deviation to allow for comparison across realms. Solid lines indicate median differences ± 90% confidence interval. Intervals overlapping the dotted line indicate a lack of statistically significant differences at α =0.10. b) Differences for each 2-Ma time bin.

**Fig. 3.**
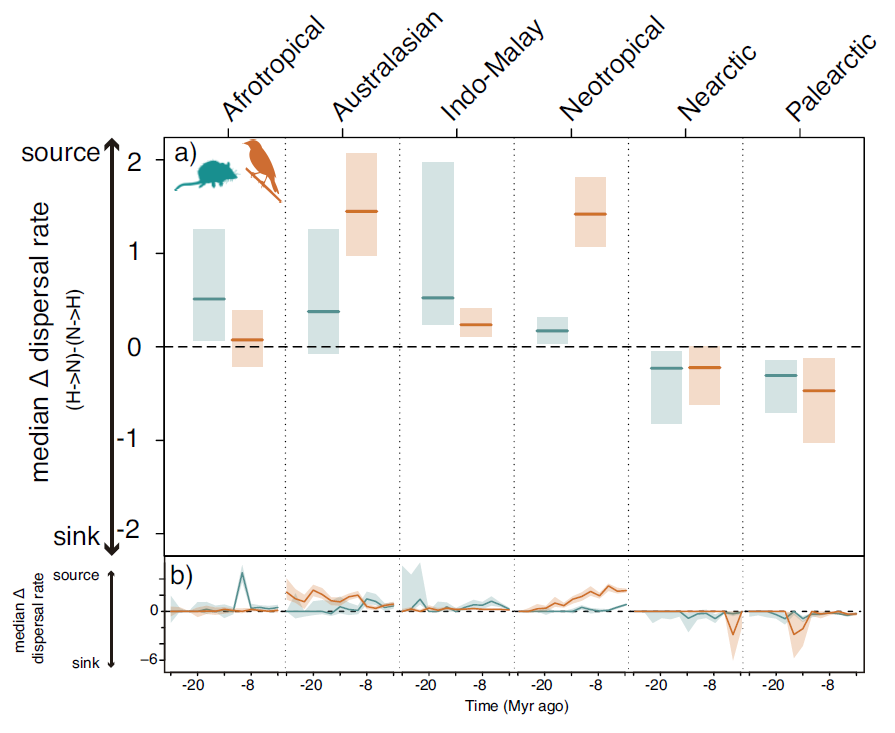
Source-sink dynamics of hotspots and their surrounding regions. a) Dispersal rates between 2 to 26 Ma ago from non-hotspots to hotspots (N→H) were subtracted from hotspot to non-hotspots (H→N) rates within each realm. Lines and shaded areas presented as in Figure 2.

Together, our findings suggest that contrasting macroevolutionary routes have shaped the uneven distribution of biodiversity across biogeographic realms. In all primarily tropical realms, except the Afrotropics, hotspots consistently generated and exported species at higher rates than their nearby areas, whereas the disproportionate richness in hotspots of temperate realms could be explained by greater rates of immigration from surrounding regions. The Afrotropics may lay somewhere outside these two routes. Afrotropical hotspots did not generate species more quickly or import them at greater rates in the last 25 Ma. The region became more arid during the late Miocene and early Pliocene as the Sahara Desert was formed (*29*). This change in the regional climate could have generated differences in the extinction dynamics of the Afrotropics hotspots compared to the non-hotspots. However, we could not estimate such extinction dynamics with the available methods. Hotspot diversity may have also been greater for ecological rather than evolutionary reasons, e.g. greater niche space (*15*).

Our results were generally robust to the methodological assumptions. First, species richness is positively correlated with region size (*30*), but we found no evidence that the difference in size between hotspots and non-hotspot regions could alone explain our results. As hotspots of endemicity were defined globally, there were large differences between the sizes of the hotspots and non-hotspot regions within each realm (Table S2). To assess whether these differences could generate the different macroevolutionary patterns that we observed, we repeated our analysis by randomly sampling combinations of cells with similar size and spatial structure to the hotspot cells in each realm. We consistently found that the real estimates of *in situ* cladogenesis (Figs. S5-S6), dispersal (Figs. S7-S8) and colonization time (Fig. S9) lay outside the simulated distributions. Therefore, the differences in macroevolutionary patterns that we observed between hotspots and their surrounding areas must have stemmed from differences in species composition and/or environmental features rather than simply due to size. Second, we also found that hotspots were more clustered in space than non-hotspot cells across all realms, particularly in tropical as compared with temperate realms. However, the differences across realms were small and <10% in the most extreme case (Table S3, Fig. S10). The slightly greater clustering of tropical hotspots is therefore unlikely to explain fully the different macroevolutionary routes that we found in tropical and temperate realms (Fig. S10). Third, we confirmed that two alternative ways of delineating biodiversity hotspots were congruent with the results for the WE-based hotspots (Supplementary Text, Figs. S11-S14). These alternate definitions used species richness (Fig. S15) and areas where narrow-ranged species occurred (Fig. S16), which have been proposed to reflect past opportunities for speciation (*31*).

Finally, we found evidence that unique environments inside the hotspots could have promoted differences in macroevolution when compared to neighboring non-hotspot regions. Linear models with spatial autocorrelation allowed us to compare environmental features potentially associated with differences in rates of *in situ* diversification and dispersal between hotspots and their surrounding areas within each realm. We found that hotspots had a greater mean net primary productivity, terrain ruggedness index and more habitats than their surrounding regions (Fig. 4). These differences were consistent across realms despite the contrasting macroevolutionary routes between tropical and temperate regions. This finding is not entirely surprising because the same environmental features can be associated with contrasting macroevolutionary routes. Specifically, increased historic opportunities for speciation may have resulted from higher energy availability (*14*) and spatial complexity (*17, 18*) in the tropics. In temperate regions, the same variables may have elevated carrying capacities and packed more immigrant species into the hotspots, which could have also acted as biodiversity refuges during past climate change (*32, 33*). The extent of the tropics was also larger over much over the past 25 Ma (*3*). Thus, more immigrants could have dispersed into temperate latitudes, i.e. “out-oftropics” hypothesis (*34*), providing an additional explanation for the different macroevolutionary routes that we observed across realms.

**Fig. 4.**
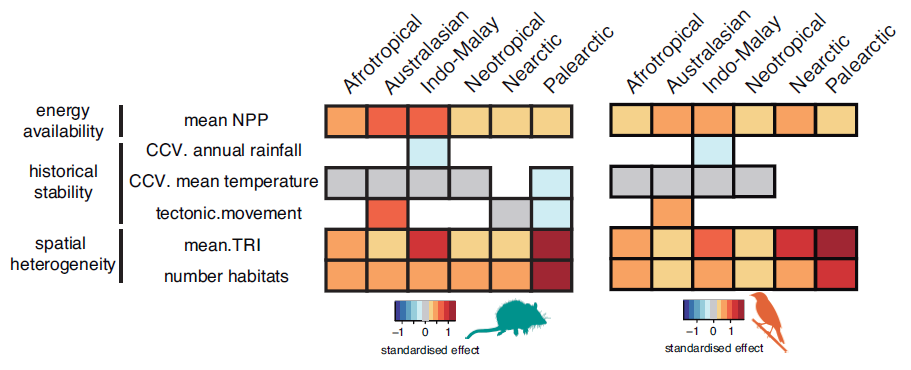
Hotspots are more spatially complex and have more energy than surrounding regions. Standardized differences in the average cell values of environmental variables between hotspots and non-hotspot cells within each of 6 biogeographic realms based on the coefficients of univariate spatially-autocorrelated linear regressions. Blank areas indicate non-significant differences. NPP = net primary productivity; CCV = climate change velocity; TRI = terrain ruggedness index.

Our study offers an integrative approach to understand why biodiversity varies so much across the globe by using a regional-scale and spatially-explicit reconstruction of historical dispersal and diversification alongside an analysis of ecological gradients. Generally, vertebrate, invertebrate and plant diversity are spatially correlated at regional scales across the planet (*4*), so we expect similar mechanisms to generate biodiversity across the Tree of Life (*35*). Similar analyses carried out in other groups may nevertheless result in clade-specific idiosyncrasies. For instance, the relative roles of *in situ* cladogenesis and dispersal as drivers of regional diversity may be different in taxa with lower vagility than mammals and birds, such as amphibians and insects (*36, 37*). By simultaneously comparing different macroevolutionary routes and their macroecological features in two major vertebrate clades, our study now provides a new answer to the old question of why diversity varies so much across the world.

## Acknowledgments

We thank T. Jucker, V. Soria-Carrasco and J. Garcia-Porta for useful comments that helped improve the manuscript. We thank the Gatsby Charitable Foundation (Grant Number GAT2962), Wellcome Trust (Grant Number 105602/Z/14/Z) and Isaac Newton Trust (Grant Number 17.24r) for funding. J.I and A.J.T designed the study. J.I. performed the analysis. J.I. and A.J.T interpreted the analysis and wrote the manuscript. The authors declare no competing interests. Relevant scripts will be uploaded to github at github.com/javierigea/xxxx

**List of Supplementary Materials:**

Materials and Methods

Figs. S1 to S18

Tables S1 to S5

References (38-64)

Supplementary Text

